# Inactivation of Influenza A Viruses (H1N1, H5N1) During Grana-Type Raw Milk Cheesemaking: Implications for Foodborne Transmission Risk

**DOI:** 10.1101/2025.06.18.660327

**Authors:** Ana Moreno, Stefano Pongolini, Giuseppe Merialdi, Giovanni Cattoli, Calogero Terregino, Nicola Santini, Stefano Benedetti, Luisa Loli Piccolomini, Anna Padovani, Alfonso Rosamilia, Giovanni Loris Alborali, Paolo Daminelli

## Abstract

**Background:** The detection of H5N1 highly pathogenic avian influenza virus (HPAIV) in lactating dairy cattle in the United States, with high viral titers in raw milk, has raised concerns about potential zoonotic transmission through the consumption of unpasteurized milk and raw-milk dairy products. While inactivation studies exist for pasteurized milk, data on virus persistence during the manufacture of raw-milk cheeses remain scarce.

**Aim:** To evaluate the survival and inactivation of avian influenza viruses (AIV), including both low pathogenic (LPAIV, H1N1) and highly pathogenic (HPAIV, H5N1) strains, during the production and ripening of Grana-type hard cheeses made from raw cow’s milk.

**Methods:** Experimental cheesemaking was conducted using raw milk artificially contaminated with A/duck/Italy/281904-2/06 (H1N1; 107.75 EID_50_/mL) or A/duck/Italy/326224-2/22 (H5N1 clade 2.3.4.4b; 106.75 EID_50_/mL). Cheeses were produced in accordance with Parmigiano Reggiano production standards and ripened for 30 days at 5–6 °C. Viral presence was assessed in finished cheeses by inoculation on SPF embryonated chicken eggs (ECE), hemagglutination (HA) assay, and monoclonal antibody-based ELISA.

**Results:** No infectious virus was detected in any cheese sample produced from contaminated milk following two blind passages in SPF-ECE. Both HA and ELISA tests yielded negative results, indicating complete inactivation of the virus.

**Conclusion:** This study demonstrates that the traditional Grana-type cheese production process— including curd cooking, acidification, and ripening—effectively inactivates both LPAIV and HPAIV, even at high contamination levels. These findings support the microbiological safety of hard cheeses made from raw milk with regard to AIV, contributing to risk assessment and food safety policies during avian influenza outbreaks.

## Introduction

Highly pathogenic avian influenza (HPAI) continues to pose a substantial threat to animal and public health globally, particularly affecting poultry and wild birds [1]. Among the various HPAI subtypes, H5N1 influenza A virus (HPAIV) remains of particular concern due to its zoonotic potential and continued global spread. The H5N1 viruses currently circulating are derived from the A/Goose/Guangdong/1/1996 (GsGd) strain, first identified in China in the late 1990s. These viruses have undergone extensive reassortment, giving rise to multiple genetic variants. Since October 2020, H5N1 viruses of clade 2.3.4.4b have been detected across Europe, following reassortment between H5N8 and wild bird-origin N1 viruses. From autumn 2021 onward, clade 2.3.4.4b viruses have become dominant and have demonstrated efficient geographic spread via migratory birds. Notably, these viruses exhibit a high capacity for reassortment with co-circulating avian influenza A viruses and the ability to infect mammals, especially bird-feeding species [1]. In late 2021, clade 2.3.4.4b was reported in Canada, marking its first detection in the Americas [2]. These strains, genetically similar to European viruses, have since reassorted with low-pathogenicity influenza viruses endemic to North American avifauna, generating novel genotypes with diverse internal gene constellations.

In March 2024, the first case of H5N1 infection in cattle was reported in Texas, United States, challenging previous assumptions regarding bovine resistance to influenza A viruses [3,4]. Subsequent investigations revealed widespread infection: by early 2025, 1,048 confirmed outbreaks had been reported in dairy herds across 17 U.S. states [5]. Whole genome sequencing confirmed these viruses as clade 2.3.4.4b, genotype B3.13, containing four gene segments of North American origin (NS, PB1, PB2, NP) and four of Eurasian origin (HA, NA, MA, PA) [3,6]. Clinical signs reported in H5N1-infected cattle have included respiratory and gastrointestinal symptoms, decreased milk production, and altered milk appearance—often colostrum-like—along with dehydration and lethargy [6,7]. Of these, changes in milk quality and production volume are the most readily observed, as they are detected during routine milking. Tests on raw milk samples from symptomatic animals revealed, quite unexpectedly, extremely high viral titers—reaching up to 108 TCID_50_/mL—and a strong affinity of the virus for mammary gland tissue [8]. These unusually high viral loads, together with evidence of lethal H5N1 infections in farm cats that consumed raw milk from infected cows, have raised urgent concerns about the safety of the U.S. dairy supply.

Following the surge in outbreaks among dairy cattle and poultry, several human cases have been reported in the U.S., primarily among agricultural workers, with 70 confirmed infections and one death [9]. Although 2024 saw widespread detection of H5N1 in birds and mammals across nearly all global regions [10], the total number of human cases remains relatively low (n = 85) [11]. Importantly, approximately 80% of these cases occurred in the U.S.—the only country to report widespread infections in lactating cattle—underscoring the need to assess the potential for bovine-to-human transmission [12].

Although U.S. regulations mandate the disposal of abnormal milk prior to pasteurisation [13,14], asymptomatic or preclinical infections could result in contaminated milk entering bulk storage. A study conducted in 2024 detected influenza A virus RNA in 20.2% of pasteurised milk and dairy samples via quantitative real-time RT-PCR (qRT-PCR), though no infectious virus was recovered, supporting the efficacy of pasteurisation in inactivating the virus [15]. Nevertheless, the presence of high viral loads in raw milk and potential exposure through unpasteurised dairy products remain public health concerns.

Hard cheeses produced from raw milk, such as Grana-type cheeses, are of particular interest. These cheeses—exemplified by Parmigiano Reggiano and Grana Padano—are widely consumed and exported. Their traditional manufacturing process involves raw milk, natural skimming, the addition of natural whey, curd cooking, and extended ageing. To date, limited data exist on the behaviour and inactivation of influenza A viruses during traditional cheesemaking, especially in European-style products.

The aim of this study was to investigate the persistence and inactivation of two influenza A subtypes (H1N1 and H5N1) during the production of Grana-type cheeses. The cheesemaking process was simulated in a controlled laboratory setting, reproducing key steps including raw milk usage, natural whey inoculation, curd cooking, and storage under whey. Preliminary trials were conducted to validate virus detection and isolation methods in cheese matrices and to assess virus stability in ultra-high temperature (UHT) milk. These trials informed the design of the main experimental study.

## Materials and methods

### Validation of sample preparation

To assess the suitability of cheese homogenates for virus isolation and titration in specific pathogen-free embryonated chicken eggs (SPF-ECEs), a preliminary test was conducted using two 1-g portions of Grana cheese. One sample served as a negative control, while the second was inoculated with 1 mL of an H1N1 viral suspension (hemagglutination [HA] titer: 1:128 per 25 µL). Both samples were cut into small pieces using sterile scissors and transferred into 15 mL Falcon tubes containing three 4.5 mm stainless steel beads and 10 mL phosphate-buffered saline (PBS) supplemented with antibiotics (final concentration: 10% w/v homogenate).

Samples were homogenised using a TissueLyser II (Qiagen Italia, Milan, Italy) at 30 Hz for 5 minutes. From each sample, two 1 mL aliquots were prepared: one filtered through a 0.45 µm syringe filter, the other left unfiltered. All aliquots were inoculated into SPF embryonated chicken eggs (SPF-ECEs) as described in section 2.2. The matrix’s potential toxicity and its compatibility with virus detection in eggs were evaluated after 4 days of incubation at 37 °C.

### Virus isolation and titration in embryonated chicken eggs

Before inoculation, SPF-ECEs were candled to confirm embryo viability, and the air sac line was marked. The eggshell surface was disinfected with iodine tincture before and after puncturing. For virus isolation, 200 µL of each sample was inoculated into the allantoic cavity of 9–10-day-old SPF eggs, using at least four eggs per sample. Eggs were candled daily for four days. Embryos that died within 4 days were chilled at 5–6 °C for a minimum of 4–5 hours before sample collection.

At 4 days post-inoculation, allantoic fluids were harvested and tested by HA assay (1% chicken red blood cells) and monoclonal antibody-based sandwich ELISA (MAb-NPAEL) targeting influenza A nucleoprotein (ATCC HB65, H16-L10-4R5). Two blind passages were performed before confirming the absence of viral growth.

For virus titration, 10-fold serial dilutions of milk or cheese homogenates were prepared in antibiotic-supplemented PBS. Four eggs were inoculated with 100 µL of each dilution. At 4 days post-inoculation, allantoic fluid was collected from dead embryos (≥48 h) and from live embryos. The 50% egg infectious dose (EID_50_/mL) was calculated using the Reed and Muench method.

### Inactivation of influenza A virus in ultra-high temperature (UHT) milk

To assess virus inactivation during a thermal treatment simulating cheese production, UHT semi-skimmed milk was experimentally inoculated with H1N1 virus (A/Duck/Italy/281904-2/06). Two 300 mL milk batches were prepared: one contaminated with virus (kept at 4–5 °C under agitation for 10 minutes), the other serving as a negative control.

Both milk batches were subjected to a heat treatment simulating the thermal profile of Parmigiano Reggiano cheesemaking. The treatment consisted of holding the milk at 32 °C for 10 minutes, followed by 53 °C for 50 minutes. These steps were performed using two calibrated water baths maintained at the respective temperatures. Milk samples were placed in two identical glass beakers during heating, and temperature monitoring was conducted using a thermometer inserted in the control beaker to ensure accuracy.

Post-treatment, 10 mL samples were collected from the inoculated batch. Viral titers were quantified by infecting embryonated chicken eggs (ECEs) to determine the 50% egg infectious dose (EID_50_/mL), according to standard protocols. Viral titers before and after heat treatment were compared to assess inactivation efficacy (see Table 1).

**Table 1.**
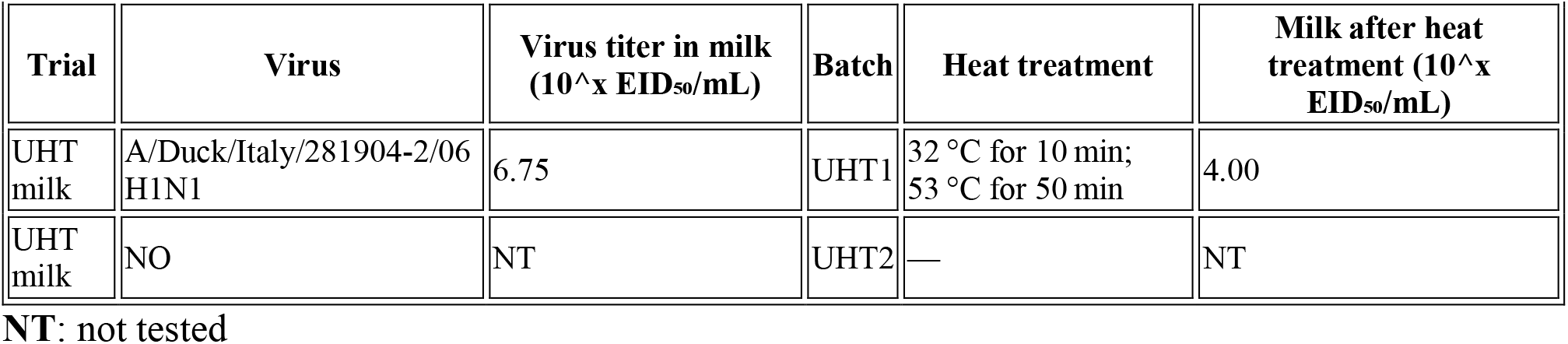
Summary of viral titration in UHT milk following heat treatment.

| Trial | Virus | Virus titer in milk (10<sup>x</sup> EID<sub>50</sub>/mL) | Batch | Heat treatment | Milk after heat treatment (10<sup>x</sup> EID<sub>50</sub>/mL) |
| --- | --- | --- | --- | --- | --- |
| UHT milk | A/Duck/Italy/281904-2/06 H1N1 | 6.75 | UHT1 | 32 °C for 10 min;<br>53 °C for 50 min | 4.00 |
| UHT milk | NO | NT | UHT2 | — | NT |

NT: not tested

### Cheese production process

The cheesemaking process was conducted to replicate the traditional Parmigiano Reggiano PDO method [16], adapted for laboratory conditions as follows:

1. **Milk Collection and Skimming:** Raw whole cow’s milk was collected and stored chilled at no less than +8 °C. Natural skimming was performed by allowing the milk to stand for 16– 18 hours at 11–12 °C, enabling fat separation.
2. **Starter Whey Addition:** Skimmed milk was supplemented with 3% (w/w) starter whey to promote acidification during subsequent steps.
3. **Coagulation:** The milk mixture was heated to 32 °C and held for 8–10 minutes to simulate the addition of rennet and the onset of curd formation.
4. **Curd Cooking:** The temperature was increased to 53.5 °C and maintained for 50 minutes to mimic curd cooking and maturation under whey.
5. **Molding and Salting:** The curd was transferred into molds and salted according to standard procedure.
6. **Ripening:** Cheeses were ripened for 30 days at 6–8 °C under controlled humidity conditions.

Samples were stored in sealed containers, and manipulations requiring opening were performed under a laminar flow hood. Curds were ripened in open molds, and internal cheese portions were used for virus testing.

### Inactivation of influenza A virus during Grana-type cheese production First trial: H1N1 virus

To assess the inactivation of influenza A virus during the production of Grana-type cheese, an initial trial was conducted using an avian H1N1 virus strain (A/Duck/Italy/281904-2/06). The protocol was applied to three 1-liter batches of raw cow’s milk (batches A, B, and C), processed simultaneously under laboratory conditions simulating Parmigiano Reggiano PDO cheesemaking steps.

Batches A and B were artificially contaminated with H1N1 virus at a known titer, while batch C served as an uncontaminated negative control.

All three batches underwent the full cheesemaking procedure:

1. Natural skimming by surfacing (16–18 h at 11–12 °C)
2. Addition of 3% starter whey
3. Heating to 32 °C for 8–10 minutes
4. Addition of rennet
5. Cooking phase at 53.5 °C for 50 minutes
6. Molding, salting, and ripening at 6–8 °C for 30 days

For batch A, samples were collected at key stages of the production process:

- Starter milk
- Skimmed milk and cream fraction (post natural surfacing)
- After addition of starter whey (post 32 °C pre-rennet phase)

In all collected samples, viral titers were determined by endpoint dilution assay in specific-pathogen-free embryonated chicken eggs (SPF-ECEs), and expressed as EID_50_/mL.

After ripening, three internal 1 g portions were collected from each cheese produced in batches A and B (total of six samples). Each sample was tested undiluted by inoculation into SPF-ECEs for viral growth detection. All eggs underwent two blind passages and were evaluated by hemagglutination assay (HA) and monoclonal antibody immunoassay (MAb-NPAEL).

**Figure 1.**
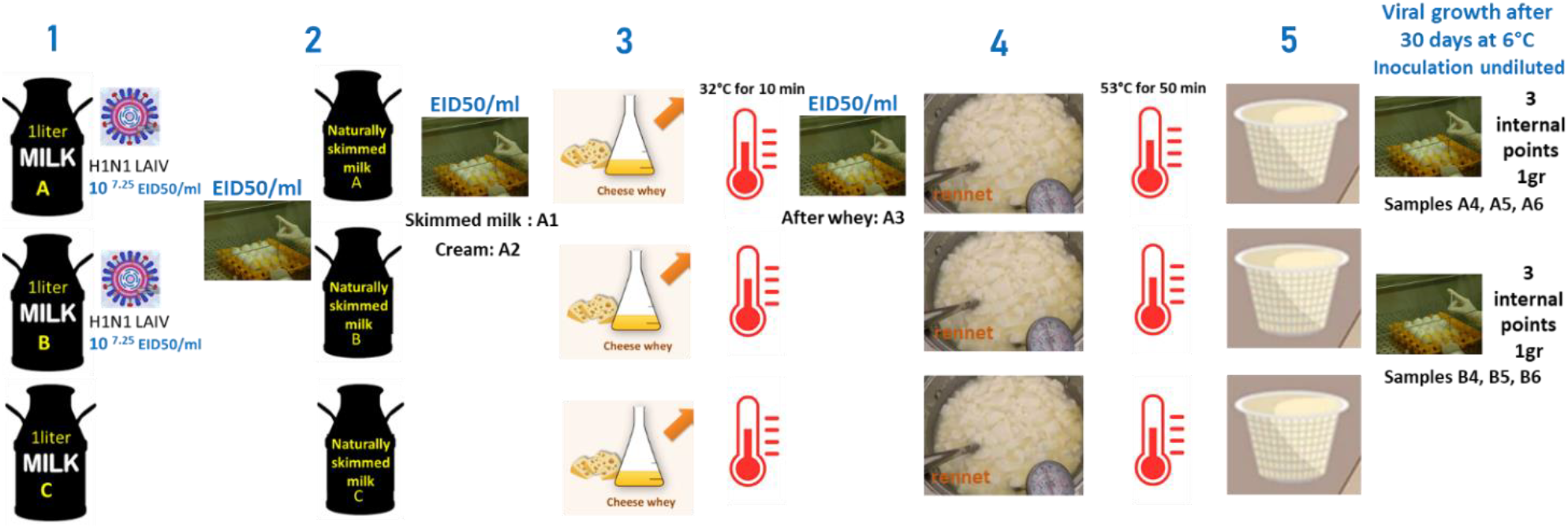
Trial 1 workflow: 1-Inoculation of 2 L raw milk with H1N1 virus; preparation of batches A and B (1 L each), and batch C (control); 2-Skimming; testing of skimmed milk (A1) and cream (A2); 3-Addition of starter whey; sample A3 collected post-heating; 4-Coagulation, curd cooking (53.5 °C, 50 min), and whey maturation; 5-Moulding, salting, and ripening; collection of three internal 1 g samples per cheese (A4–A6, B4–B6).

### Second trial: H5N1 virus

A second trial was conducted using a high-pathogenic avian H5N1 virus (A/duck/Italy/326224-2/22, clade 2.3.4.4b) to confirm inactivation under identical cheesemaking conditions. Two 1-liter milk batches (D and E) were processed in parallel in BSL-3 conditions.

Viral titration was performed on initial milk, and finished cheeses were tested after 30 days of ripening. Three internal 1 g samples per cheese were collected and tested using SPF-ECEs and confirmatory HA and MAb-NPAEL assays. The experimental setup was analogous to the H1N1 trial (see Figure 3).

**Figure 2.**
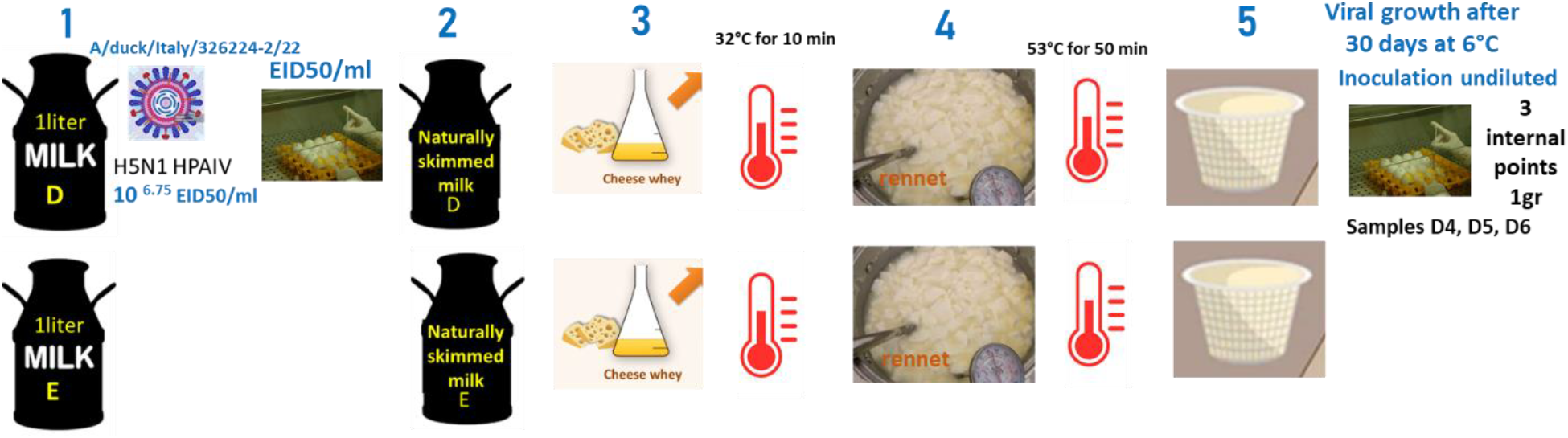
Trial 2 workflow: 1-Inoculation of 1 L raw milk with H5N1 virus; preparation of batch D and E (control); 2-Skimming; 3-Addition of starter whey; 4-Coagulation, curd cooking (53.5 °C, 50 min), and whey maturation; 5-Moulding, salting, and ripening; collection of three internal 1 g samples per cheese (D4-D6).

## Results

### Preliminary trials

The preliminary validation of the dairy matrix confirmed both the absence of toxic effects on SPF-ECEs and the ability to support viral growth. The sample inoculated with H1N1 showed a higher HA titer after filtration through a 0.45 μm filter (mean: 1:512 per 25 μL) compared to the unfiltered sample (mean: 1:192 per 25 μL), based on two replicates. Consequently, filtered samples were used for subsequent viral growth assays, while unfiltered 1:10 dilutions were used for titration in SPF-ECEs.

Inactivation tests of the H1N1 virus in UHT semi-skimmed milk revealed a reduction in viral titer exceeding 2 log10 units. However, complete inactivation was not achieved, requiring the subsequent cheese production trials (Trials 1 and 2) to evaluate virus survival following the full cheesemaking and ripening process. Results are summarized in **Table 1**.

### First trial with low-pathogenic H1N1 virus

In Trial 1, two liters of raw milk were artificially contaminated with the H1N1 virus (titer: 10^7.25 EID_50_/mL) and divided into two batches (A and B). Only batch A underwent viral titration at different processing steps. Titrations in skimmed milk (Sample A1), cream (Sample A2), and milk post-starter whey addition (Sample A3) revealed reductions of less than 2 log10 units. Due to ethical considerations regarding animal use, titrations were performed from dilutions −1 to −6, sufficient for an approximate quantification of viral decrease.

After 30 days of ripening at 6–8 °C, samples from cheeses produced in batches A and B were tested. No viral growth was detected in any of the six internal 1 g cheese samples after two blind passages in SPF-ECEs, as confirmed by negative results in HA assay and MAb-NPAEL. **Table 2** summarizes these findings.

**Table 2.** Summary of virus titration and growth assays during cheese production trials.

| Trial | Virus | Virus titer in milk (10<sup>x</sup> EID<sub>50</sub>/mL) | Batch | Skimmed milk | Cream | After starter whey | Cheese after 30 days ripening (1 g samples) |
| --- | --- | --- | --- | --- | --- | --- | --- |
|  |  |  |  | A1 | A2 | A3 | A4-A5-A6 |
| 1 | A/Duck/Italy/281904-2/06 (H1N1) | 7.25 | A | >5.50 | >5.50 | 6.00 | NEG |
|  |  | 7.25 | B | NT | NT | NT | NEG |
|  | NO | NO | C | NT | NT | NT | NT |
| 2 | A/Duck/Italy/326224-2/22 (H5N1) | 6.75 | D | NT | NT | NT | D4-D5-D6<br>NEG |
|  | NO | NO | E | NT | NT | NT | NT |

NT: not tested; NEG: no viral growth detected after two blind passages.

### Second trial with high-pathogenic H5N1 virus

Trial 2 confirmed the findings of Trial 1. Milk contaminated with HPAIV H5N1 (clade 2.3.4.4b) at a titer of 10^6.75 EID_50_/mL was used to produce one 1 L cheese batch (D) under BSL-3 conditions. Batch E served as an uncontaminated negative control. After 30 days of ripening, all internal cheese samples tested negative for virus replication, again confirmed by two blind passages with negative HA and MAb-NPAEL results. Due to biosafety constraints and the objective of evaluating final virus inactivation, no intermediate titrations were performed during processing (see **Table 2**). Given the absence of viral replication at 30 days, no additional ripening or sampling was performed.

### Chemical and physical properties of the cheeses

The raw milk used in this study had a total bacterial count of 12,000 CFU/mL and 311,000 somatic cells/mL, consistent with typical dairy production in Italy’s Po Valley. Its chemical composition was representative of national standards, with fat content of 4.2 g/100 g, protein at 3.6 g/100 g, and casein at 2.85 g/100 g. Following natural skimming, the fat-to-casein ratio was adjusted to between 0.80 and 1.05, in accordance with the Parmigiano Reggiano PDO specification. The starter whey (3% w/w), with an acidity of 31 °SH/50 mL, was sourced from a Grana Padano dairy and was used to promote proper acidification—an essential step for curd formation and subsequent maturation under whey.

Given the small volume of milk processed and, consequently, the limited amount of curd and cheese obtained, chemical-physical characterization was assessed using the Moisture on Fat-Free Basis (MFFB), following the criteria established by Commission Decision 97/80/EC [17]. This parameter is considered more representative than pH or water activity (Aw) alone when evaluating whether a product conforms to the properties of Grana-type cheeses.

According to European classification standards, cheeses are categorized based on MFFB values as follows:

Soft cheese: MFFB ≥ 68%

Semi-soft cheese: 62% ≤ MFFB < 68%

Semi-hard cheese: 55% ≤ MFFB < 62%

Hard cheese: 47% ≤ MFFB < 55%

Extra-hard cheese: MFFB < 47%

The experimental protocol yielded cheeses with MFFB values between 47% and 55% after approximately 4 weeks of ripening. This places the products within the “hard cheese” category and is consistent with typical Grana-type cheeses such as Parmigiano Reggiano and Grana Padano, which normally undergo 9 months or more of maturation. The 30-day ripening period at 4 °C used in this study represents a conservative simulation compared to traditional PDO cheesemaking, which typically occurs at higher temperatures and over longer durations (9–12 months). Importantly, water activity in Grana-type cheeses generally declines to below 0.93 by the end of minimum ripening, contributing to an environment that is increasingly hostile to viral survival.

The successful inactivation of virus observed in this study should be attributed to the combined effect of critical processing steps, including natural skimming, acidification via starter whey, curd cooking and maturation under whey, salting, and extended ripening. Together, these steps form a robust multibarrier system that ensures the microbiological safety of the cheese—even in the absence of thermal pasteurization.

## Discussion

### Foodborne Transmission Risk of HPAIV

The global dissemination of H5N1 HPAIV, particularly widespread infection of lactating cattle in the United States, has raised urgent concerns regarding the potential for zoonotic transmission through dairy products.

While human infections were previously attributed primarily to direct contact with infected birds, mammals, or contaminated environments, the detection of high viral loads in milk from infected cattle highlights the need to assess foodborne transmission risks—especially via consumption of unpasteurized milk and derived products.

Key risk factors include: the infectious viral load in milk; virus stability under refrigeration; efficacy of processing steps in reducing infectivity; and host susceptibility, particularly the minimal infectious dose [18]. H5N1 has been shown to preferentially target bovine mammary tissue, where both avian-type (α2,3-linked) and human-type (α2,6-linked) sialic acid receptors are expressed, facilitating high-level viral shedding into milk [8,19].

Furthermore, the virus can remain viable in refrigerated milk (4 °C) for over five weeks [20], underscoring the importance of downstream processing in mitigating transmission risk.

### Impact of Cheesemaking Processes on Virus Inactivation

Implementing effective milk treatments and production processes is critical to ensuring the inactivation of viruses before dairy products are consumed. Pasteurization remains the gold standard for inactivating avian influenza viruses (AIVs) in milk and dairy products, as consistently demonstrated in previous studies [20–23]. While viral RNA has occasionally been detected in thermally treated products such as butter, cheese, and ice cream, no viable virus has been recovered, confirming the efficacy of standard pasteurization protocols in inactivating AIVs. However, the fate of highly pathogenic avian influenza viruses (HPAIVs) during the manufacture of raw-milk cheeses has been largely unexplored [24].

Our study addressed this critical knowledge gap by experimentally assessing the inactivation of both low-pathogenic (H1N1) and highly pathogenic (H5N1 clade 2.3.4.4b) influenza A viruses during the production of Grana-type hard cheese from raw cow’s milk under conditions conforming to Parmigiano Reggiano PDO specifications. Cheese samples were analyzed in triplicate after 30 days of ripening to assess residual viral infectivity. In both experimental arms, no viable virus was detected in any sample, indicating complete inactivation under these production conditions. Given this consistent outcome, the ripening period was not extended.

This outcome is especially relevant given that the initial viral titers employed in our experiments (10^7.25 EID_50_/mL for H1N1 and 10^6.75 EID_50_/mL for H5N1) exceeded both the mean and maximum viral loads reported in naturally contaminated milk samples from recent U.S. outbreaks (mean: 10^3.5 EID_50_/mL; max: 10^6.3 EID_50_/mL) [20]. Furthermore, regulatory frameworks mandate the exclusion of visibly altered milk from processing, further minimizing the potential risk of consumer exposure. These considerations strengthen the relevance and applicability of our findings.

The Production Regulations of Parmigiano Reggiano PDO establish a stringent framework of rules and standards that must be followed by all authorized dairies throughout the entire cheesemaking process. Only producers affiliated with the Parmigiano Reggiano Consortium— founded in 1934—are permitted to manufacture this iconic Italian cheese under the PDO label. The Consortium is tasked with the protection, oversight, enhancement, promotion, and international representation of Parmigiano Reggiano, ensuring both the quality of the product and the integrity of its production process. By defining both the technological procedures and the compositional requirements of the final product, the Consortium guarantees uniformity across all certified producers. This standardization minimizes variability and ensures that cheeses labeled as Parmigiano Reggiano consistently meet the high safety and quality benchmarks established by the PDO specification.

### Comparison with Literature

In the H1N1 trial, intermediate titration after skimming, cream separation, and whey addition revealed ∼1 log10 reduction in viral load, confirming that these steps alone do not fully inactivate the virus. This aligns with prior reports that moderate heat (e.g. 50–54 °C) is insufficient to ensure complete viral inactivation [20–22].

While pasteurization remains the gold standard for eliminating viral pathogens in milk, our findings support that the integrated raw-milk cheesemaking process—including curd cooking above 52 °C, whey fermentation, and extended ripening—can similarly achieve complete inactivation of AIV. Our results are consistent with those of Nooruzzaman et al. [24], who reported that viable H5N1 could be recovered from raw-milk cheeses with pH ≥5.8, but not from those acidified to pH 5.0 after 60 days of ripening. To date, that study is the only other investigation of H5N1 survival in raw-milk cheese. The final pH in Parmiggiano Reggiano cheese (∼5.3) was closer to this inactivation threshold, underscoring the critical role of acidification in virus control.

Although precise pH and water activity (Aw) values were not measured in our trials due to sample constraints (low amount of curd/cheese, not representative of the real shape and size of an Italian “grana-type” PDO cheese), the calculated MFFB value confirmed the final product was well representative of the authentic Parmigiano Reggiano PDO cheese. In PDO-compliant production, both pH and Aw progressively decline during ripening, creating unfavorable conditions for viral persistence.

Finally, our experiments did not extend to the full 9-month ripening period required for Grana PDO cheeses. However, given that complete inactivation occurred by day 30, and that further decreases in pH and Aw are expected during aging, it is highly unlikely that viable virus would persist in cheese ripened in accordance with PDO specifications.

## Conclusions

This study provides robust experimental evidence that Grana-type hard cheeses produced from raw milk using traditional methods—as defined by the Parmigiano Reggiano PDO specification— effectively inactivate both low-pathogenic (H1N1) and highly pathogenic (H5N1 clade 2.3.4.4b) avian influenza viruses.

Despite the use of viral inocula exceeding titers reported during natural outbreaks, no viable virus was recovered from cheese samples after 30 days of ripening. This inactivation is attributable to the cumulative impact of key technological steps, including milk skimming, starter culture addition, curd cooking, whey removal, salting, and ripening.

Importantly, all procedures adhered strictly to PDO specifications, reinforcing the reproducibility and real-world relevance of the findings. These results strongly support the microbiological safety of raw-milk Grana-type cheeses, even in the event of contamination with highly pathogenic AIV strains.

When traditional processes are properly applied, the risk of foodborne HPAIV transmission through raw-milk hard cheeses is negligible. These findings also highlight the critical role of processing parameters—especially temperature, acidification, and ripening duration—in ensuring viral inactivation and protecting public health.

## Statements

## Ethical statement

No ethical approval was necessary for this study.

## Data availability

None

## Acknowledgements

We thank Ms Francesca Adella (IZSLER) for their technical support.

## Authors’ contributions

Study design and conceptualisation: AM, PD, GM, SP, CT, SB, NS

Material collection: AM, PD

Laboratory diagnostics: AM, PD

Writing—original draft preparation: AM, PD, AR

Manuscript revision: AM, PD, SP, GLA, GC, NS, SB, LLP, AP

Commenting and editing: all authors

